# Barley Long Non-Coding RNAs and Their Tissue-Specific Co-expression Pattern with Coding-Transcripts

**DOI:** 10.1101/229559

**Authors:** Gökhan Karakülah, Turgay Unver

## Abstract

Long non-coding RNAs (lncRNA) with non-protein or small peptide-coding potential transcripts are emerging regulatory molecules. With the advent of next-generation sequencing technologies and novel bioinformatics tools, a tremendous number of lncRNAs has been identified in several plant species. Recent reports demonstrated roles of plant lncRNAs such as development and environmental response. Here, we reported a genome-wide discovery of ~8,000 barley lncRNAs and measured their expression pattern upon excessive boron (B) treatment. According to the tissue-based comparison, leaves have a greater number of B-responsive differentially expressed lncRNAs than the root. Functional annotation of the coding transcripts, which were co-expressed with lncRNAs, revealed that molecular function of the ion transport, establishment of localization, and response to stimulus significantly enriched only in the leaf. On the other hand, 32 barley endogenous target mimics (eTM) as lncRNAs, which potentially decoy the transcriptional suppression activity of 18 miRNAs, were obtained. Presented data including identification, expression measurement, and functional characterization of barley lncRNAs suggest that B-stress response might also be regulated by lncRNA expression via cooperative interaction of miRNA-eTM-coding target transcript modules.

## 1. Introduction

Long non-coding RNAs (lncRNA) are known as non-protein coding transcripts longer than 200 nt. Although, the information about their functions is so limited, studies revealed that they have several direct and indirect roles in the transcriptional, post-transcriptional or post-translational processes such as gene expression, chromatin modification, transcriptional regulation, and conformational changes in proteins (reviewed by Liu et al., 2015). They act as the precursor of micro-RNA (miRNA) and short interfering-RNA (siRNA). For instance, five lncRNA in Arabidopsis (npc34, npc351, npc375, npc520, and npc523) matched with 24-nt siRNAs from both strands, suggesting these lncRNAs are siRNA precursor. Moreover, it was reported that plant miRNAs, miR869a and miR160c, which were derived from lncRNAs of npc83 and npc521, respectively (Ben Amor et al., 2009). Additionally, it was also discovered that miR675 is derived from a mouse lncRNA, H19, and extensively expressed in embryonic liver (Dey et al., 2014; Keniry et al., 2012).

Recently, short peptide-coding sequences were discovered in the non-coding regions of plant primary-miRNAs (pri-miRNA) and called as miRNA-encoded peptide (miPEP), which increases the transcription of pri-miRNA (Lauressergues et al., 2015; Waterhouse and Hellens, 2015). Therefore, it was suggested that some plant lncRNAs might have peptide coding potential (Liu et al. 2015). In addition, miRNA activity can be regulated by endogeneous target mimicry (eTM) molecules, being a type of lncRNAs (Karakulah et al., 2016). Such an example, an endogenous lncRNA called *Induced by Phosphate Starvation 1 (IPS1)* of *Arabidopsis thaliana* binds to miR399 to inhibit the cleavage of the miR399 target transcript (Franco-Zorrilla et al., 2007). Other than these functions, plant lncRNAs were found to be involved in many regulatory mechanisms, such as histone modeling (Heo and Sung, 2011), promoter modification (Ding et al., 2012; Zhou et al., 2012), protein re-localization (Sousa et al., 2001), and alternative splicing (Bardou et al., 2014).

To date, lncRNAs have been identified extensively in mammals, in which human genome includes more than 56,000, and mice have almost 46,000 lncRNAs (Xie et al.,2014). Publicly available databases such as LncRNAdb (http://lncrnadb.com), a database for functional lncRNAs, harbor functionally annotated lncRNAs, of the majority belong to the human. A few databases were so far released for plant lncRNAs; such as the Green Non-Coding Database (GreeNC, http://greenc.sciencedesigners.com) (Paytuvi Gallart et al., 2016) and CANTATAdb (http://cantata.amu.edu.pl) (Szczesniak et al., 2016), which provides information for around 45,117 lncRNAs from several plant species. RNA-sequencing or deep transcriptome analysis is an important technology, which provides information not only for protein coding transcripts, but also for noncoding RNAs (such as miRNA, siRNA, piwi-interacting RNA (piRNA), and small-nucleolar RNA (snoRNA) as well as lncRNAs). In addition, it allows distinguishing lncRNAs expressed in different tissues or cells.

Boron (B) is an essential micronutrient for plants, and its unfavorable concentration negatively affects plant growth and productivity where the soils having with insufficient or excess B (Unver et al., 2008). The range of sufficient B concentration in soil is so limited, thus generally plants suffer from either B-deficiency or -toxicity problem that is common in agricultural soils of around the world (Camacho-Cristobal et al., 2015). However, plants growing in B-contaminated soils must tolerate the excess level of B to survive. In the last decade, many studies conducted to understand the cellular mechanisms underlying to balance cellular B content in plants (Miwa and Fujiwara, 2010; Tombuloglu et al., 2015). Facilitated transport of boric acid (a regular form of B in soil) by transporter channels was suggested to be the molecular regulatory mechanism. Several B-importer and exporter proteins have recently been identified as B-transporters to regulate its cellular homeostasis (Miwa and Fujiwara, 2010). On the other hand, novel sequencing-based approaches to discover the transcriptional response at genome-wide level are being extensively utilized in plants faced with unfavorable environmental conditions. In this context, to quantify gene expression and to annotate coding-transcripts, we performed a high-throughput genome-wide transcriptome analysis on barley tissues treated with excess B (1 mM), previously (Tombuloglu et al., 2015). In addition, we also screened B-responsive miRNA expression pattern (Ozhuner et al., 2013); identified MYB type transcription factors (TF) (Tombuloglu et al., 2013) and water channel Aquaporins (AQP) (Tombuloglu et al., 2016) to understand B homeostasis of barley. These studies helped to observe the main or possible players involved in B regulation. Besides the emerging evidences suggested that the molecular regulatory mechanism is so complicated and not only limited to activity of those coding transcripts. LncRNAs being the new players were also be discovered as regulatory molecules on the regulation of gene expressions.

To date, a large set of RNA-seq libraries was used to identify lncRNAs in genome-wide scale or tissue/condition/inoculation-specific manner (Chen et al., 2016; Li et al., 2014; Liu et al., 2012). Stress-responsive lncRNAs were examined from the RNA-seq data of the plants under the distinct type of stress conditions as well (reviewed by Chekanova, 2015; Shafiq et al., 2016; Zhang et al., 2013). Qi et al., 2013 identified 584 lncRNAs, which were responsive to drought stress in foxtail millet. 125 putative lncRNAs were identified in wheat, responsive to powdery mildew infection and heat stress (Xin et al., 2011). Detailed examination of a large set of poplar (*Populus trichocarpa*) RNA-seq data revealed 504 lncRNAs in response to drought (Shuai et al., 2014). Additionally, Huang et al reported over 12,000 barley lncRNAs, of them 604 were *Fusarium* head blight inoculation responsive (Huang et al., 2016). However, no such a study to profile expression level of lncRNAs under the B-excess as one of the abiotic stress conditions was conducted till now. In this study, we identified and quantitatively compared the expression of B-responsive barley lncRNAs from four transcriptome datasets. Tissue-specific (root and leaf) and excess B-responsive lncRNAs, which were co-expressed with coding-transcripts, discovered and comprehensively analyzed.

## 2. Materials and Methods

### 2.1. Identification of barley lncRNAs

To study global expression profiling of barley lncRNAs under boron stress condition, we utilized four transcriptome libraries each including pooled RNAs from tree biological replicate previously generated by our group (SUB337351 and SUB2170217) (Tombuloglu et al., 2015). Briefly, a total of 208,249,690 clean sequencing reads from four paired-end libraries (50_leaf; 52, 422,032, 50_root; 52,168,358, 1000_leaf; 52,305,062, and 1000_root, 51,354,238) were utilized in this study. We first removed the adapter sequences and low-quality reads from the sequencing reads with Trimmomatic v0.36 (Bolger et al., 2014), and these clean reads were aligned to the barley reference genome (ASM32608v1 assembly) by TopHat2 v2.1.1 with default parameters (Kim et al., 2013). Afterward, genome-aligned reads were assembled *ab initio* using popular transcriptome analysis suit Cufflinks v2.2.1 (Trapnell et al., 2010) to build potential transcript structures. All gene transfer format (GTF) files produced in the assembly step were merged with Cuffmerge tool, and transcript features were queried against the Ensembl Plant database (release 33) (Kersey et al., 2016) by Cuffcompare to discover previously unannotated transcript sequences. Among all unannotated transcripts, including coding and non-coding sequences, we obtained lncRNAs as follows: (i) we first tested coding potential of each transcript individually using TransDecoder (https://transdecoder.github.io/), and filtered out those with an open reading frame having more than 100 amino acids, (ii) we then removed transcripts which were shorter than 200 nucleotides in length, (iii) to remove housekeeping lncRNA species, we queried all potential lncRNAs against to non-coding RNA family database, Rfam (v12.1) (Nawrocki et al., 2015), with Infernal tool (v1.1.1; cutoff Evalue ≤ 0.001) (Nawrocki et al., 2009). Then, potential miRNA precursors, tRNAs etc were removed and were not included for further analysis, and (iv) we aligned all potential lncRNA transcripts to the Swiss-Prot database (release 2017_01) (The UniProt, 2017) using Blastx (v2.5.0; cutoff E-value ≤ 0.001) (Camacho et al., 2009) to eliminate transcripts with potential protein-coding ability.

### 2.2. Expression pattern analysis of coding and non-coding RNAs

Transcript abundances in each library were measured with Kallisto v0.43.0 (Bray et al., 2016). Then the transcripts expressed <1 <u>T</u>ranscripts <u>P</u>er <u>M</u>illion (TPM) in all libraries were considered as transcriptional noise and were removed from further downstream analysis steps. Differentially expressed transcripts within each group (leaf and root samples) were determined by calculating fold changes of TPM values in RNA-seq datasets. Transcripts with differential expression values ≥ 2 fold-changes in compared datasets were classified as boron responsive. The Gene Ontology (GO) analysis of differentially expressed genes was performed using online GO analysis toolkit, agriGO applying default parameters (Du et al., 2010).

### 2.3. Co-expression analysis of lncRNAs with coding mRNAs and prediction of endogenous target mimicry (eTM) sequences

We predicted putative functions of differentially expressed lncRNAs with “guilt-by-association” approach, which employed in previous studies for lncRNA annotation (Rinn and Chang, 2012) (Guo et al., 2013) (D’Haene et al., 2016). To reveal potential lncRNA-mRNA associations, we identified co-expressed mRNA-lncRNA pairs with Spearman’s correlation test in R v3.1.0 statistical computation environment (Team, 2016). Then, co-localized mRNA-lncRNA pairs on the reference genome were identified with Bedtools v2.25.0 (Quinlan, 2014). We considered the mRNA-lncRNA pair as co-expressed if the Spearmen’s rho is equal or greater than 0.90 (p-val<0.01) between the expression values of coding and lncRNA transcripts, and as co-localized when the distance between two transcripts were less than 10 kb..To dissect putative eTM sequences among the transcripts annotated as lncRNA, we employed our analysis pipeline previously introduced by our group (Karakulah et al., 2016). In the eTM sequences analysis pipeline, we utilized mature miRNA sequences of barley collected from miRBase (release 21) (Griffiths-Jones, 2006). To identify potential target transcripts of barley miRNAs, we utilized psRNATarget, an online miRNA target analysis tool (Dai and Zhao, 2011) as previously described previously (Akdogan et al., 2016; Bakir et al., 2016; Eldem et al., 2012; Inal et al., 2014; Yanik et al., 2013).

## 3. Results and discussion

### 3.1. Barley lncRNA identification

After the adapter sequence trimming and removal of low-quality reads, the mean library size of four sequencing libraries included in the study was over 20 million (min: 18223418, max: 22171430). Additionally, we observed an average of 82.35 % (min: 78.7 %, max: 90.9 %) overall read mapping for the alignment step, and considered all sequencing libraries had sufficient quality to perform an *ab initio* transcriptome reconstruction analysis. The run of Cufflinks pipeline was led to the identification of as many as 34,000 previously unannotated intergenic transcripts, of which 10,439 were the lack of coding potential and more than 200 bp in length (Table S1). When we filtered out the transcripts expressed at low levels (< 1 TPM in all samples), we obtained 8,009 intergenic putative lncRNAs in the final list, which were distributed almost equally to all barley chromosomes (Figure 1A). However, chromosome 2 was observed to be the richest one in terms of the total number of lncRNAs it harbors. In this study, a total number of ~8,000 barley lncRNAs were identified which is smaller than that of human (~56,000) (Xie et al., 2014) and mouse (~46,000); higher than fruit fly (~3,300) (Chen et al., 2016), and poplar (2,542) (Shuai et al., 2014) (Table S1). Actual numbers of lncRNAs can be altered depending on sample examined. In this analysis, four RNA-Seq libraries were used to detect total lncRNAs. More lncRNAs can be found from barley genome by increasing the number of RNA-Seq sets from distinct tissues and/or conditions. In general, low expression levels of most lncRNAs compared to protein-coding genes make it more difficult to detect lncRNAs (Mercer et al., 2011). Generally, they are excluded from the total lncRNA pool resulting fluctuations of total lncRNA number. But it is important to note that lncRNAs with low expression may have a big impact, thus extensive and a deep pipeline is required to extract lncRNA, which may possess important biological functions. In general, the distribution of lncRNAs to the barley chromosomes is proportional with its chromosome sizes, except chromosome 2, which includes the highest number (Fig1A).

**Figure 1.**
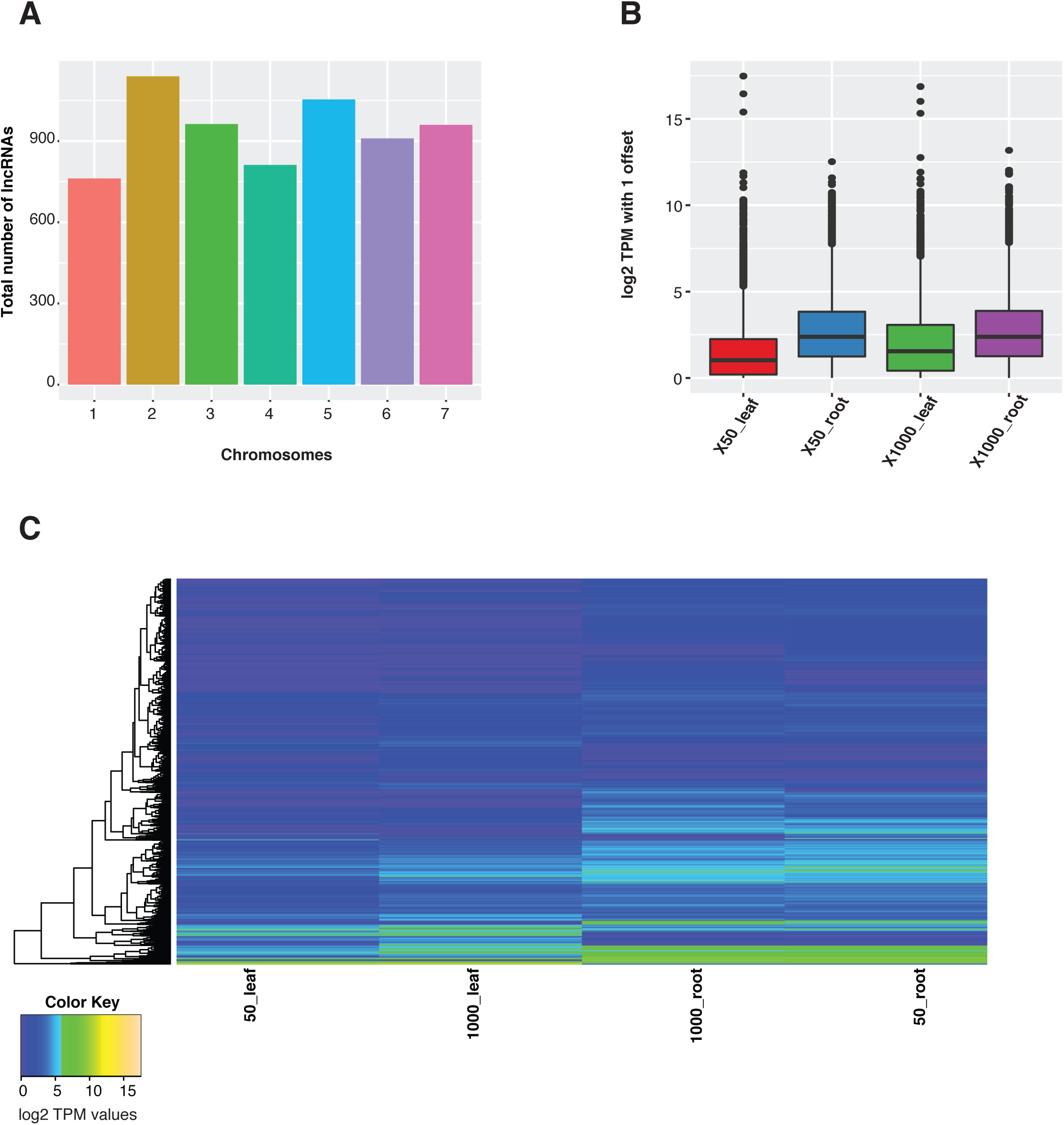
Chromosomal distribution of barley lncRNAs and their expression in normal and stress conditions. A. Total number of lncRNAs localized to each barley chromosome (from chromosome 1 to 7). B. Box plot representation of expression levels of lncRNAs before and after boron exposure in root and leaves. C. Expressional changes of lncRNAs across samples are being illustrated as heat map graph.

### 3.2. Expression pattern of barley lncRNAs and coding transcripts upon excess B-treatment

As the expression profiles of lncRNAs in root and leaf samples were examined, it was determined that expression levels of lncRNAs in the samples collected from same tissue were similar to one another (Figure 1B). The hierarchical clustering analysis, however, revealed that particular lncRNA clusters were expressed at relatively higher levels specific to tissue types (Figure 1C). Differential expression analysis of both coding and lncRNA transcripts showed that there was 2 fold or more change (up- or down- regulation) in the expression of the vast amount of transcripts in response to boron stress in leaves and root tissues (Table S2). We observed that the number of common coding transcripts that were differentially regulated in both tissues was 517; in addition, the total number of differentially expressed coding transcripts in the leaf tissue was roughly doubled as compared to root tissue (Figure 2A). Similar to the differential expression analysis of coding transcripts, we detected a greater number of up- or downregulated lncRNAs in the leaves than the root samples (Figure 2B).

**Figure 2.**
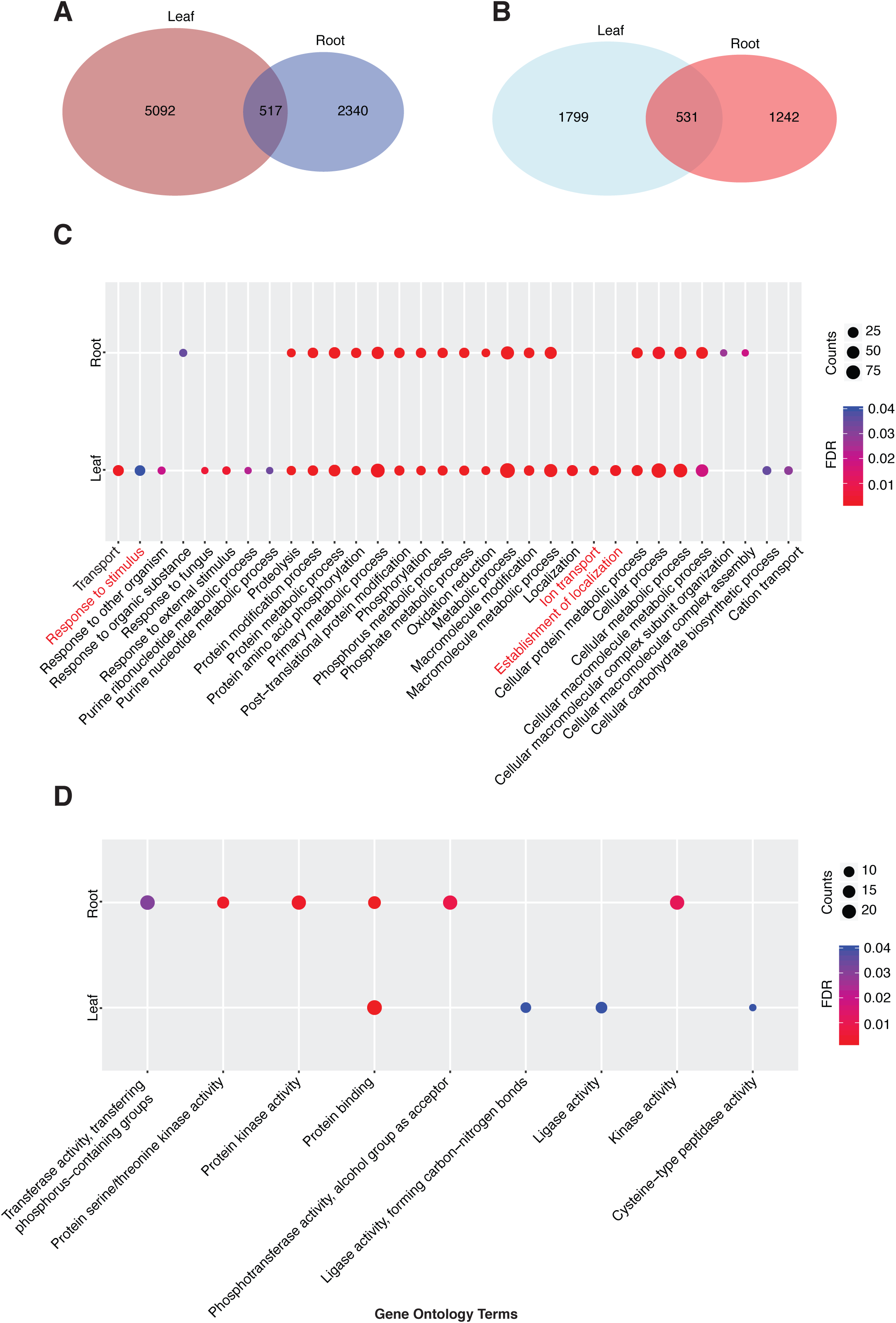
Differentially expressed transcripts and GO term enrichment analysis. A-B. Venn diagram of differentially expressed coding (A) and lncRNA (B) transcripts in leaf and root. C-D. Significant biological process (C) and molecular function (D) GO terms associated with differentially expressed coding transcripts.

### 3.3. Functional annotation of lncRNAs co-expressed with coding transcripts

The GO enrichment analysis of differentially expressed coding transcripts (Table S3) revealed that ion transport (GO:0006811), establishment of localization (GO:0051234), and response to stimulus (GO:0050896) terms significantly enriched (FDR<0.05) only in the leaf samples (Figure 2C). In addition to this, molecular function terms of ligase activity (GO:0016874) and cysteine-type peptidase activity (GO:0008234) were significant and specific to the leaves (Figure 2D). Based on the co-expression and colocalization analysis of barley lncRNAs and coding transcripts, we observed potential lncRNAs in association with the ion transports, establishment of localization, and response to stimulus related transcripts (Figures 3 and 4). We determined 6 lncRNAs, which strongly linked to ion transport related genes (Figure 3A). However, only one lncRNA (TCONS_00061958) was showing ≥ 2 fold expressional change in both the leaf and root samples (Figure 3B). A great majority of response to stimulus related lncRNAs increased (≥ 2 fold) their expression in leaf samples upon the boron exposure (Figure 3C-D).

**Figure 3.**
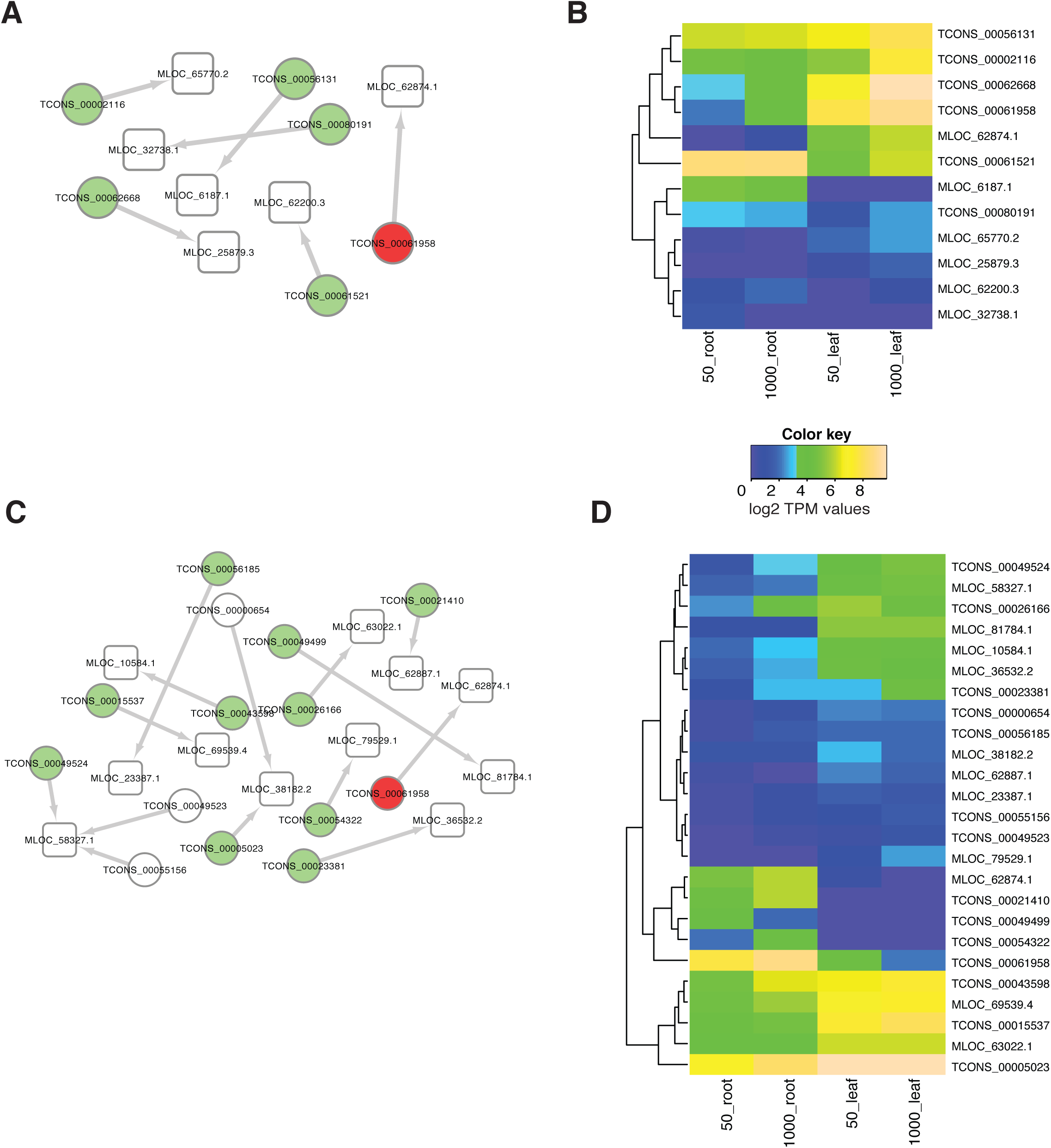
Ion transport and response to stimulus associated lncRNAs. A-C. LncRNAs and coding transcripts related to ion transport (A) and response to stimulus (C) GO terms. LncRNAs and coding transcripts are represented as circles and round rectangles, respectively. Differentially expressed lncRNAs before and after boron exposure are shown as colored. Green colored lncRNAs are differentially expressed only in leaves. However, lcnRNAs are colored as red if they are differentially regulated in both root and leaves. LncRNAs and associated coding transcripts (co-expressed and co-localized) are linked to each other with network edges. B-D. Heat map representation of expression levels both lncRNAs and coding transcripts associated with ion transport (B) and response to stimulus (D) GO terms.

**Figure 4.**
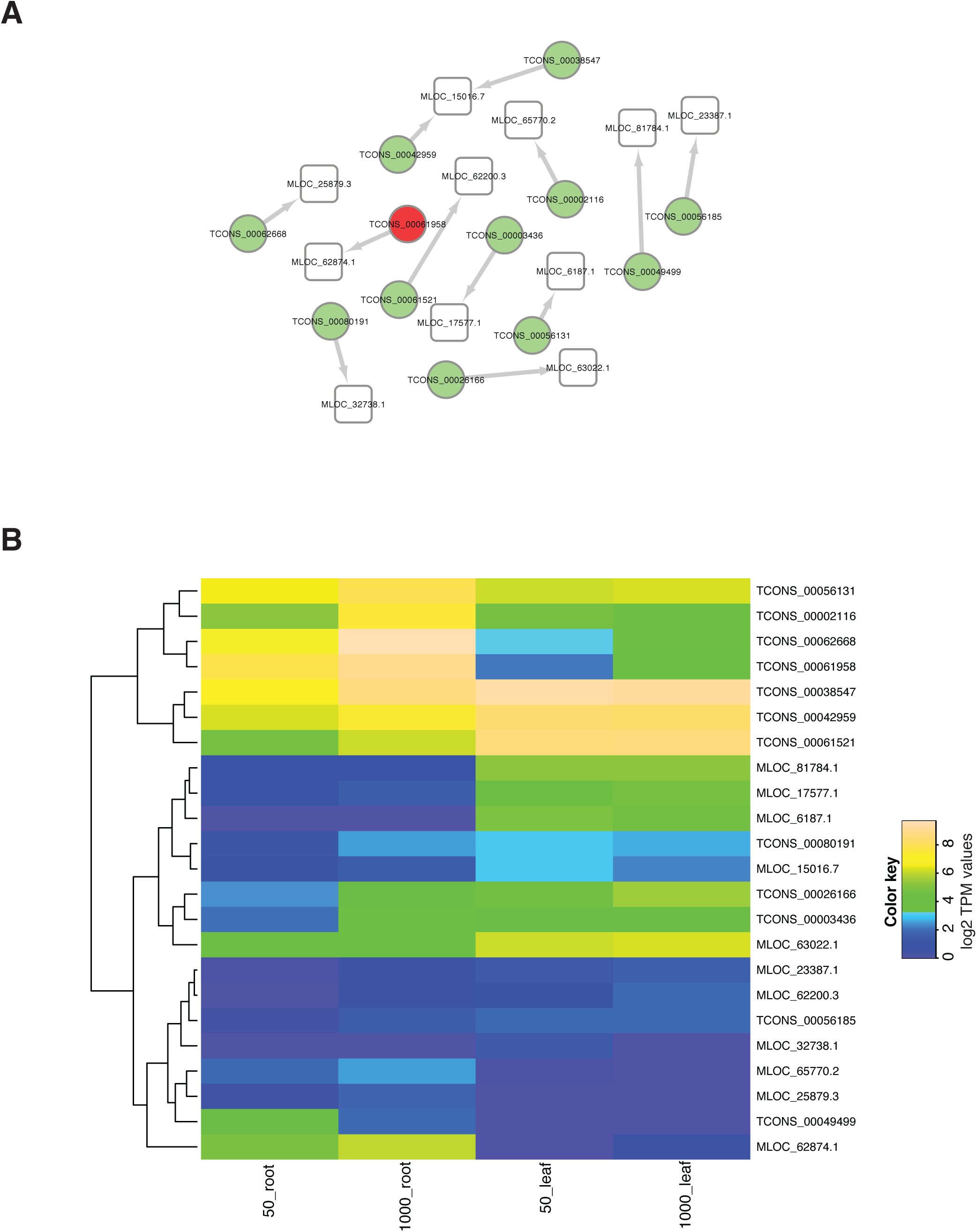
Establishment of localization associated lncRNAs. A. Network representation of establishment of localization related coding transcripts and lncRNAs (the same color scheme as in Figure 3 is used). B. Expression levels of establishment of localization GO term associated coding and lncRNAs.

Additionally, we detected 12 lncRNA sequences, which had similar expression patterns and co-localized with “establishment of localization” related genes (Figure 4). We also detected that some of the lncRNAs were only differentially expressed in tissue-specific manner such as TCONS_00002116 expression in leaf under B-excess (Figure 4B). However, TCONS_00061958 was found to be differentially regulated in both leaf and root tissues. Studies also revealed the differential expression pattern of plant lncRNAs such as Arabidopsis, wheat, and poplar in response to biotic and abiotic stresses (Liu et al., 2012; Shuai et al., 2014; Xin et al., 2011). Here, we identified that barley boron-responsive lncRNAs are expressed in the tissue-specific manner (Figures 2-4 and Table S2). According to GO term enrichment analysis, differentially expressed coding transcripts were categorized into three biological processes: response to stimulus (50 transcripts), ion transport (17 transcripts), and establishment of localization (52 transcripts), which were leaf specific in this analysis (Fig 2C and Table S3). These findings were consistent with our previous report (Tombuloglu et al., 2015), where the biological process, ion transport, and establishment of localization categories were found to be enriched in both leaf and root tissues.

### 3.4. eTM sequence discovery

Here, we predicted 32 barley eTMs, which might decoy the transcriptional suppression activity of 18 miRNAs, including conserved barley miRNAs such as miR159a, miR166a, and miR399 (Table 1 and Table S4). In the GO enrichment analysis of miRNA target genes using the agriGO tool, we found 102 significant (FDR<0.05) different GO terms in three domains, consisting of molecular function, biological process, and cellular component (Table S5). Among the terms, the most significant one was “protein amino acid phosphorylation” term (GO:0006468, FDR= 4.70E-58). Our findings in eTM analysis suggest that barley lncRNAs might regulate several distinct cellular and molecular processes via mimicking specific miRNA target transcripts. As it was previously reported by recent studies, lncRNAs might behave as a mimicry-transcript that targeted by miRNAs and fate it to degradation (Juan et al., 2013). It was firstly reported in Arabidopsis, over-expression of non-coding gene *IPS1* suppressed the miR399 expression that resulted in elevated expression of the miR399 target (Franco-Zorrilla et al., 2007). On the other hand, we determined and measured the boron-responsive barley miRNAs (Ozhuner et al., 2013). Accordingly, miR5049 was down-regulated in B-stressed leaf (three times than control leaf). Also, miR399 was over-expressed in leaf and suppressed in root tissue upon B-exposure (three times of each tissue than that of control). In this study, miR5049 and miR399 were also found to be as regulated miRNAs upon B-treatment by eTM analysis (Table S4) where TCONS_00032652 and TCONS_00043651 have predicted as the target mimic sequences, which are able to decoy the miRNA activities, respectively. Thus, expression of transcripts targeted by these miRNAs might be altered due to differential expressions of lncRNAs. *Phosphate transporter 2 (PHO2)* and *putative ubiquitin-conjugating enzyme (UBC)* were found to be the target genes of miR399. Also, *tubby protein-like* transcript was determined as the miR5049 target. In the transcriptome analysis, ubiquitin carboxyl-terminal hydrolase gene was highly up-regulated in leaf tissue upon excess B treatment (Tombuloglu et al., 2015). Interestingly, expression profiles of miRNA and its corresponding lncRNA target mimic transcript provide insights into the regulation of B-stress in plants. For instance, lncRNA TCONS_00043651, a potential target mimic sequence of miR399, up-regulated in roots (three times than that of control) upon B-exposure. Oppositely, miR399 expression was reflected with the same pattern in a negative direction (three times down-regulated). Similarly, five times increase of lncRNA TCONS_00043651 in leaf tissues may prevent the expression of miR399, which was up-regulated only three times (Table S2). These preliminary results suggest that boron regulation can be cooperatively controlled by the interaction of miRNA-eTM (lncRNA)- coding target transcript modules.

## 4. Conclusion

With the development of next-generation sequencing technologies and advancement in bioinformatics, more transcriptional datasets were generated including the units with little or no protein-coding potential. In recent years, the lncRNAs considered as regulatory molecules in several bioprocesses. Though a large number of lncRNA transcripts were identified in plants, no such genome-wide study was conducted for barley as an important crop. Another missing biological hypothesis is that the possible involvement of lncRNAs in boron-response mechanism. Here, we reported the genome-wide discovery of ~8,000 barley lncRNAs and measured their expression pattern upon excessive boron (B) treatment. Furthermore, we functionally annotated the coding transcripts, which are co-expressed with lncRNAs and showed that cooperative interaction of miRNA-eTM (lncRNA)- coding target transcript modules might regulate the boron-response in barley.

## Acknowledgements

The authors greatly appreciate ICGEB with grant no CRP/TUR16-03.

## Author Contributions

GK and TU organized the study. GK performed analyses and TU interpreted the data. GK and TU wrote the manuscript.

## Table

Table 1. Computationally identified putative eTM sequences having potential to act as miRNA sponge.

## Supplementary Materials

Table S1. Barley previously unannotated.transcrpits.bed

Table S2. Expression of lncRNAs and coding transcripts

Table S3. GO term enrichment analysis

Table S4. eTM sequences

Table S5. GO term enrichment analysis of eTM related miRNA target genes

